# Chevreul: An R Bioconductor Package for Exploratory Analysis of Full-Length Single Cell Sequencing

**DOI:** 10.1101/2025.05.27.656486

**Authors:** Kevin Stachelek, Bhavana Bhat, David Cobrinik

## Abstract

Chevreul is an open-source R Bioconductor package and interactive R Shiny app for processing and visualization of single cell RNA sequencing (scRNA-seq) data. It differs from other scRNA-seq analysis packages in its ease of use, its capacity to analyze full-length RNA sequencing data for exon coverage and transcript isoform inference, and its support for batch correction. Chevreul enables exploratory analysis of scRNA-seq data using Bioconductor SingleCellExperiment or Seurat objects. Simple processing functions with sensible default settings enable batch integration, quality control filtering, read count normalization and transformation, dimensionality reduction, clustering at a range of resolutions, and cluster marker gene identification. Processed data can be visualized in an interactive R Shiny app with dynamically linked plots. Expression of gene or transcript features can be displayed on PCA, tSNE, and UMAP embeddings, heatmaps, or violin plots while differential expression can be evaluated with several statistical tests without extensive programming. Existing analysis tools do not provide specialized tools for isoform-level analysis or alternative splicing detection. By enabling isoform-level expression analysis for differential expression, dimensionality reduction and batch integration, Chevreul empowers researchers without prior programming experience to analyze full-length scRNA-seq data.

**Data availability:** A test dataset formatted as a SingleCellExperiment object can be found at https://github.com/cobriniklab/chevreuldata.

**Availability & Implementation:** Chevreul is implemented in R and the R package and integrated Shiny application are freely available at https://github.com/cobriniklab/chevreul.

## Statement of Need

Exploratory data analysis is an important step in single cell RNA sequencing (scRNA-seq) studies that is made more challenging by the complexity of sequencing outputs. Widely used scRNA-seq analysis toolkits including Seurat [1] for R and Scanpy [2] for python enable flexible, reproducible analysis and have codified best-practices [2,3]. However, these command-line toolkits can be challenging for biologists without prior experience with relevant programming or statistics concepts. Several interactive tools for single cell data analysis do not require such experience including iSEE, cellxgene scclustviz, and Cerebro [4–7], yet these tools are designed to analyze droplet-based sequencing methods that yield 3’- or 5’-end short-read sequences and do not take advantage of Smart-seq-based or other deep, full-length RNA sequencing datasets, which can provide greater insight into transcript isoforms and cell states.

Chevreul is an open-source R Bioconductor package for exploratory analysis of scRNA-seq data, including deep, full-length scRNA-seq data, processed in the SingleCellExperiment Bioconductor or Seurat formats. The package simplifies scRNA-seq data analysis using a set of R Shiny apps to visualize scRNA-seq datasets with interactive plots. Chevreul is distinct from other interactive tools for single cell data analysis in its orientation toward full-length sequencing data captured by Smart-seq-based cDNA synthesis followed by short-read sequencing [8], which necessitates unique analyses such as measurement of isoform-specific expression and transcript exon coverage. Chevreul is also notable for its support for batch integration without the need for a coding environment. Batch integration is a primary concern for laboratories working with rare cell types and clinical samples of limiting cell number that necessitate multiple rounds of just-in-time processing, introducing technical batch effects. Chevreul includes easy to execute batch integration and data preprocessing pipelines with sensible default settings. Using the Chevreul R Shiny app, researchers can analyze data from multiple experiments with a common framework, greatly reducing the time needed to generate publication-ready plots, novel hypotheses, and insights from scRNA-seq experiments. Chevreul incorporates automated testing using the *testthat* package [9]. It can be easily installed on a Windows, Mac, or UNIX-like operating system from the current Bioconductor release and will be maintained and updated according to biannual Bioconductor releases. The package was inspired by the color theorist Michel-Eugène Chevreul and the optical illusion of the same name [10].

## Implementation

### Processing workflow

Chevreul is an R package (actually a metapackage) for processing and integration of scRNA-seq data from cDNA end-counting or from full-length short-read or long-read protocols and an R Shiny app for easy visualization, formatting, and analysis. Chevreul installs and loads three Bioconductor packages: *chevreulProcess* for processing, *chevreulPlot* for plotting, and *chevreulShiny* for loading and interactive analysis of processed scRNA-seq data (including full-length scRNA-seq) as SingleCellExperiment objects. All functionality contained in the three constituent packages is accessible directly from Chevreul.

Chevreul enables processing of an scRNA-seq dataset starting with a SingleCellExperiment object [11] constructed from a table of cell metadata with experimental variables such as treatment status, and a raw gene-by-cell or transcript-by-cell count matrix output from common tools such as *tximport* [12], *CellRanger* [13] or *STAR* [14]. If data are derived from full-length scRNA-seq protocols, transcripts may be quantified using alignment-free methods best used with well-annotated transcriptomes (Salmon, Kallisto), alignment-based methods best used to detect novel isoforms (StringTie2), or long-read methods for use with long-read sequencing data such as Oxford Nanopore or PacBio technologies (IsoQuant) [15– 18].

Isoform identities can be specified according to any version of the three major gene annotation databases [19–21] used in upstream transcript mapping and quantitation (Fig. 1A). Reads are then imported as separate transcript and gene assays, where gene read counts are the sum of the constituent transcript counts.

**Figure 1:**
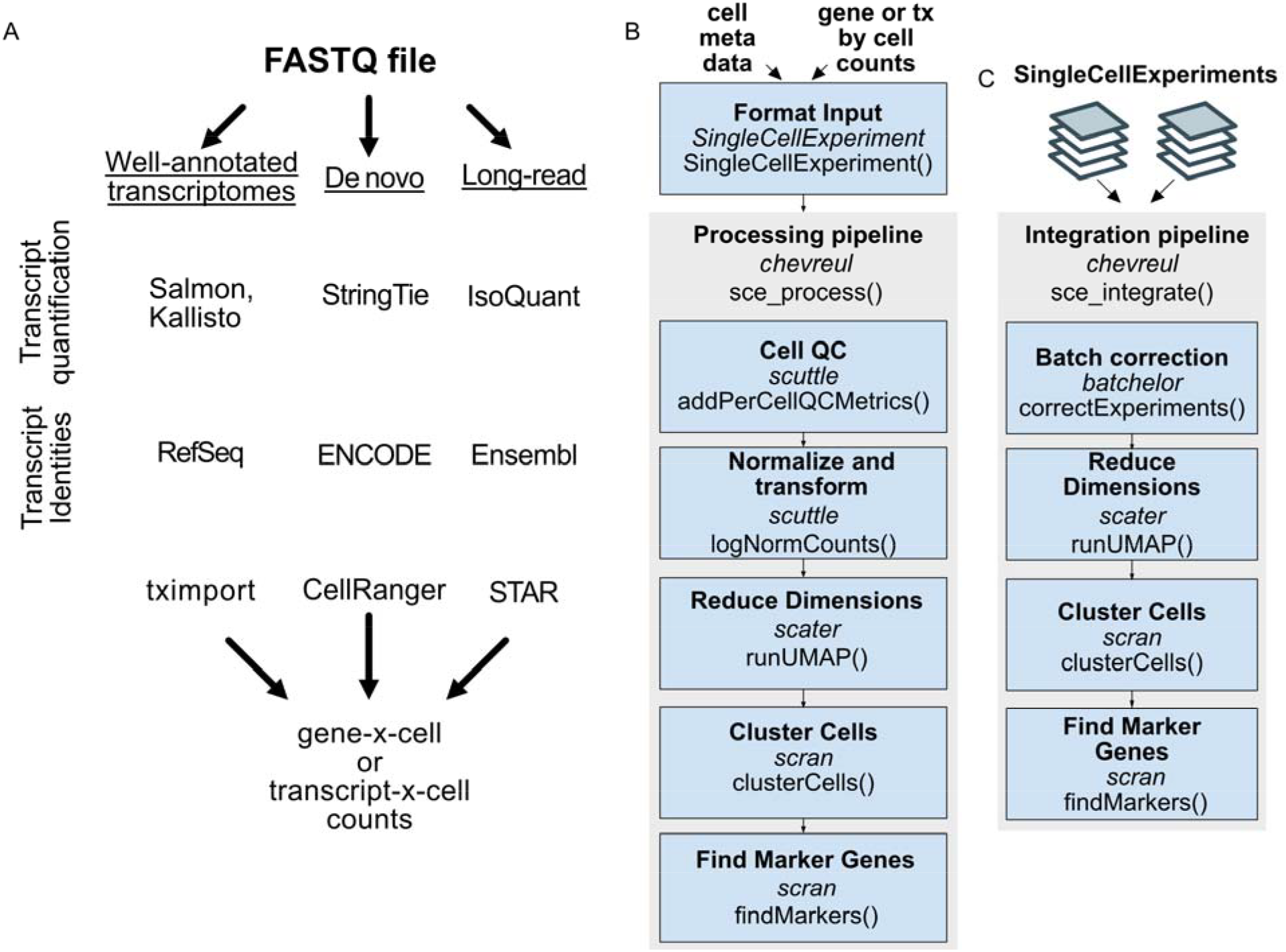
Full-length scRNAseq processing workflows and subsequent analysis with chevreul. (A) Example of workflows preceding processing and quantitation of full-length scRNA-seq datasets. (B) Processing pipeline for quality control filtering, read count normalization and transformation, dimensionality reduction, clustering, and marker gene identification. (C) Pipeline for batch integration and processing. Pipeline steps indicate *R package dependency* and specific function. See Methods for open-source software sources and Chevreul package for default settings and code.

In chevreulProcess, standardized functions prefixed with ‘sce_’ enable processing and integration of datasets. Initially, single datasets can be processed with a simple pipeline function, *sce_process()*, which allows i) quality control filtering by minimum expression and ubiquity to exclude empty droplets or degraded cells, ii) normalization by library-size scaling and log transformation, iii) dimensionality reduction by PCA, tSNE, and UMAP, iv) Louvain clustering at a range of resolutions, and v) cluster marker gene or marker transcript identification (Fig. 1B). Sensible default values are provided for all processing steps although individual steps can be adjusted at any stage. A second pipeline function, *sce_integrate()*, enables integration of a list of SingleCellExperiment objects followed by batch-corrected processing including dimensional reduction and clustering (Fig. 1C). *sce_integrate()* also implements batch correction of scRNA-seq datasets using the *batchelor* package [22]. Default settings rely on the *batchelor correctExperiments()* function to preserve the pre-existing data and metadata from input objects in the corrected output.

Chevreul enables analysis of Seurat-derived scRNA-seq objects only after their conversion to the SingleCellExperiment format. This may be preferred when a Seurat object was previously generated or when SingleCellExperiment datasets are prepared using different bioinformatic approaches or dimensional reduction parameter settings. For example, this may be desired to take advantage of alternative batch integration methods obtained using the *Seurat* package [1], which yield different normalized data input to dimension reduction and clustering compared to objects generated with the Chevreul function *sce_integrate()*. Previously generated Seurat V5 objects can be converted to SingleCellExperiment objects using the package *seuFLViz* (https://github.com/cobriniklab/seuFLViz) and further examined using Chevreul.

Recommended minimum hardware requirements for running Chevreul include 16 GB RAM. However, for larger datasets or more complex analyses, 64 GB or more is advisable. Additionally, having multiple cores can be beneficial for parallel processing. As the number of cells increases, so do the hardware requirements. For instance, a full-length scRNA-seq dataset with ∼800 cells and an average of 3.75x 10^6^ reads per cell can be analyzed with 8 GB of RAM. For larger datasets or more complex analyses, 64-128 GB of RAM can be beneficial. Runtime and memory scaling for an example dataset is reported for an ubuntu 20.04 system with 8-core Intel i7 CPU and 69 Gb RAM (Fig. 2).

**Figure 2:**
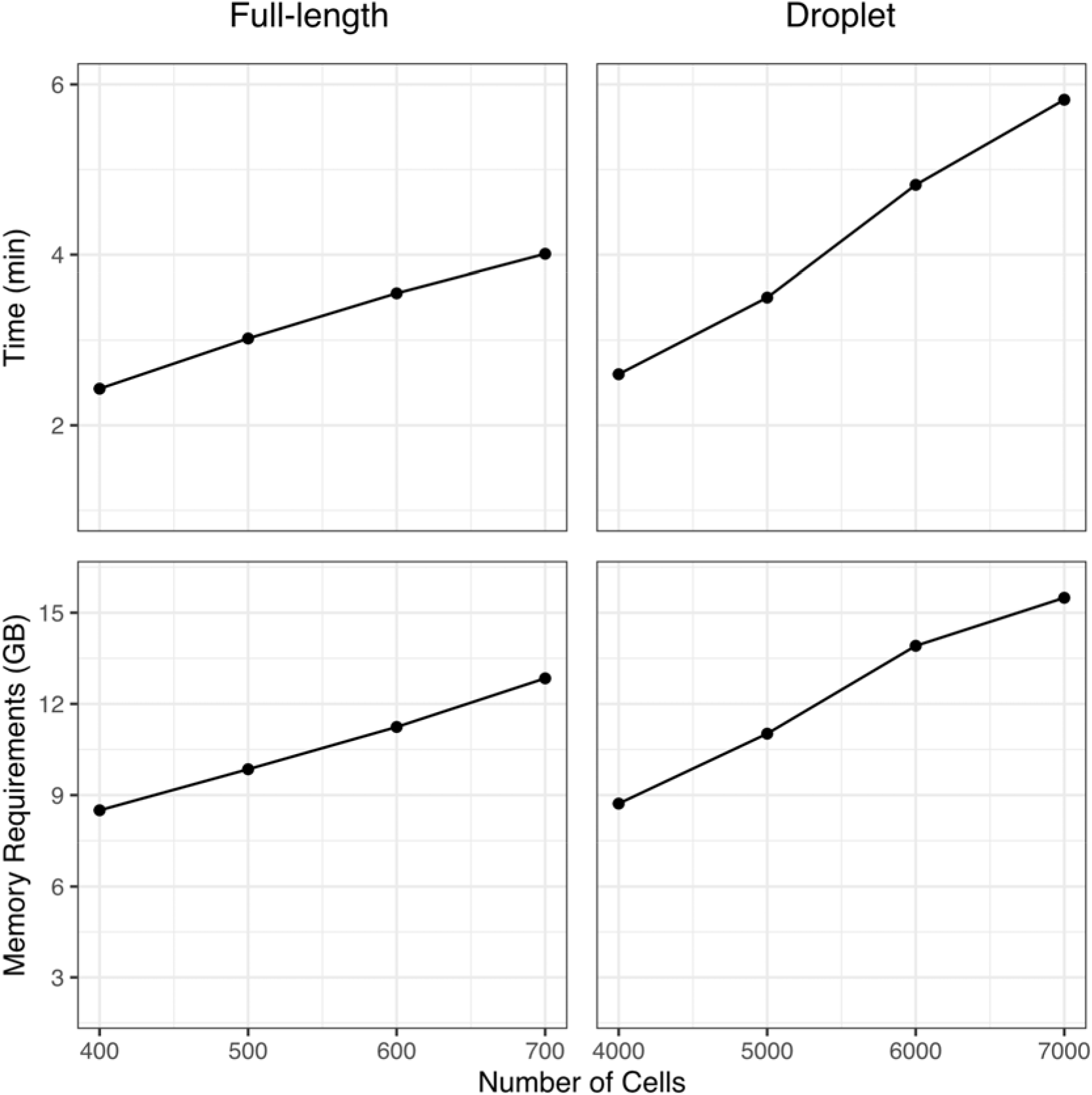
System time and memory used for initial processing of a full-length and droplet-based SingleCellExperiment. The example compares full-length scRNA-seq (794 cells, 8277 mean genes detected) and simulated droplet scRNA-seq (8000 cells, 8994 mean genes detected) datasets processed using *sce_process()* with default Chevreul settings. Genes defined using ensemble build 87. Benchmarking carried out on an ubuntu 20.04 system with 8- core Intel i7 CPU and 69 Gb RAM.

### Interactive analysis

To demonstrate the use of Chevreul, we provide a sample analysis of a publicly available Smart-seq based scRNA-seq dataset of developing human retina (GSE207802), which was previously evaluated with a Shiny app based on the *Seurat* R package [23] and is available in a preprocessed SingleCellExperiment object in the *chevreuldata* package. Chevreul documentation includes a full workflow with example vignettes including an annotated guide for the Shiny app, a walk-through of all plotting functions, and a getting-started guide for troubleshooting installation, links to relevant educational resources, background information on isoform-level analyses, and basic execution.

Starting with a processed SingleCellExperiment, the Chevreul Shiny app enables plotting of feature expression and cell metadata variables for visual analysis and data exploration. Dynamic linking between plots ensures a consistent view of a loaded dataset to uncover unrecognized relationships between experimental variables. Interactive plots are accessed on sidebar tabs on the dashboard (Fig. 3A). The Overview Plots tab allows visualization of embeddings in PCA, tSNE, or UMAP overlaid with cell metadata or feature (*i*.*e*., gene or transcript) expression (Fig. 3B). Dimensional reduction parameters such as the UMAP input dimensions and the UMAP minimum distance can be tuned to adjust embedding appearance. Alternatively, UMAP embeddings generated using Seurat can be imported into Chevreul and analyzed with the Chevreul shiny app (Supp. Fig. S1A). This can yield different, albeit similar, clusters and dimensional reduction plots due to different batch integration methods used by Seurat integration [1] and Bioconductor mnnCorrect [22], with UMAPs also affected by the selected random seed [24]. An additional section of the Overview Plots window displays read or UMI count histograms that can be colored based on metadata variables of interest such as Louvain cluster identity (Fig. 3C). Finally, a Clustering Tree plot is provided to visualize the relationship between Louvain clusters at a range of resolutions using the *clustree* package [25] (Fig. 3D).

**Figure 3:**
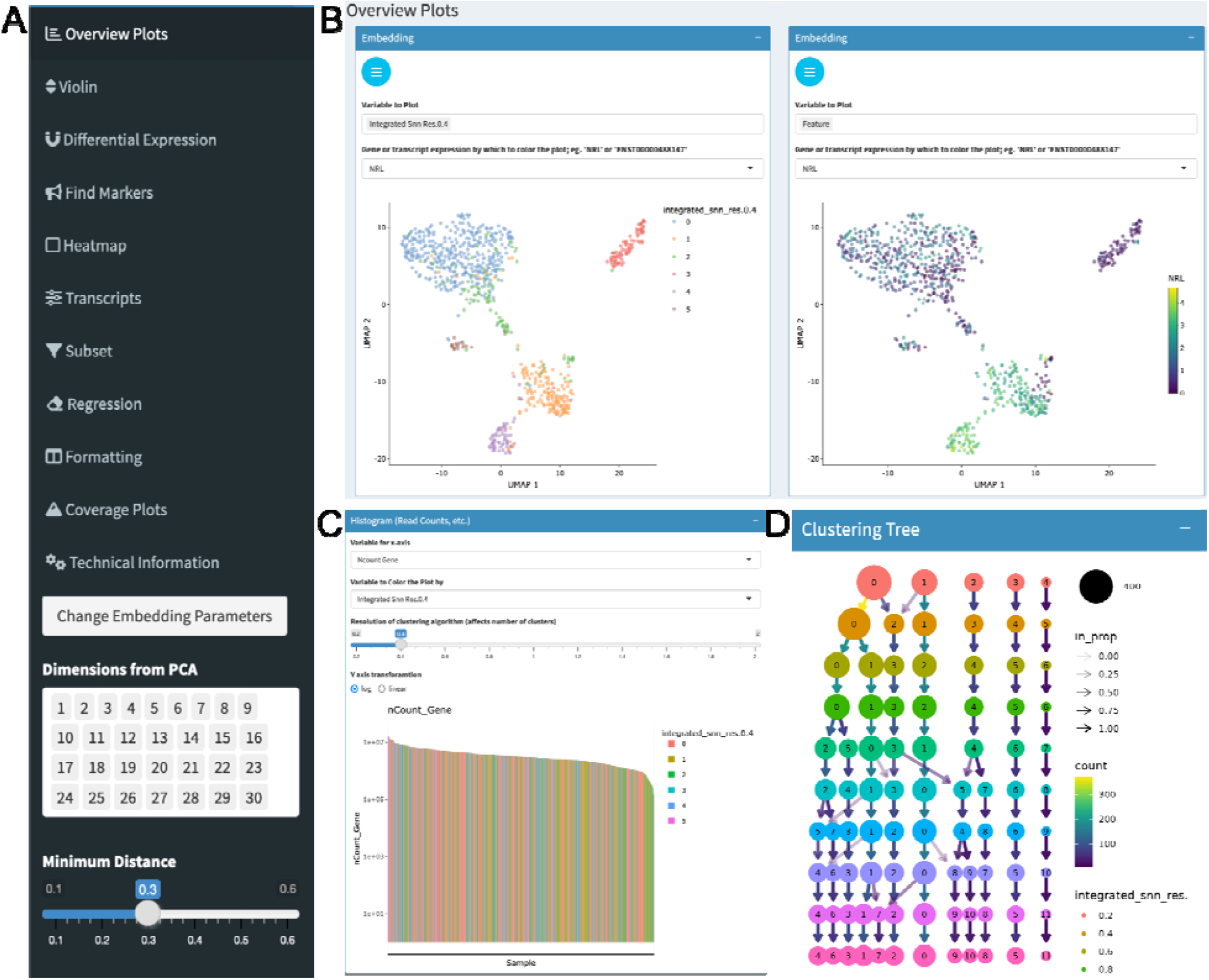
Analysis overview plots. (A) Sidebar showing Shiny app tabs. (B) UMAP embedding overlaid with metadata for cluster cell type or expression of *NRL*. (C) Read count histogram colored by cluster identity. (D) Clustering tree at multiple resolutions.

The Violin tab displays customizable violin plots of gene expression organized by cell metadata such as cell cluster, sample age, sample genotype, or multiple variables in combination (Fig. 4A). The Differential Expression tab allows differential expression analysis between groups specified by cell metadata or graphical selection from a dimensionally reduced plot. Resulting gene lists can be exported and volcano plots colored and labeled for genes surpassing desired p-values and fold-change thresholds (Fig. 4B). The Find Markers tab allows plotting of marker features such as genes or transcripts specific for cluster identities or other cell metadata and defined based on results of Wilcoxon rank-sum test. A user-assigned number of marker genes per cell group can be specified with display of those meeting adjusted p value and log fold-change thresholds (Fig. 4C). The Heatmaps tab allows plotting via complexHeatmap [26] to display selected lists of genes or transcripts (Fig. 4D).

**Figure 4:**
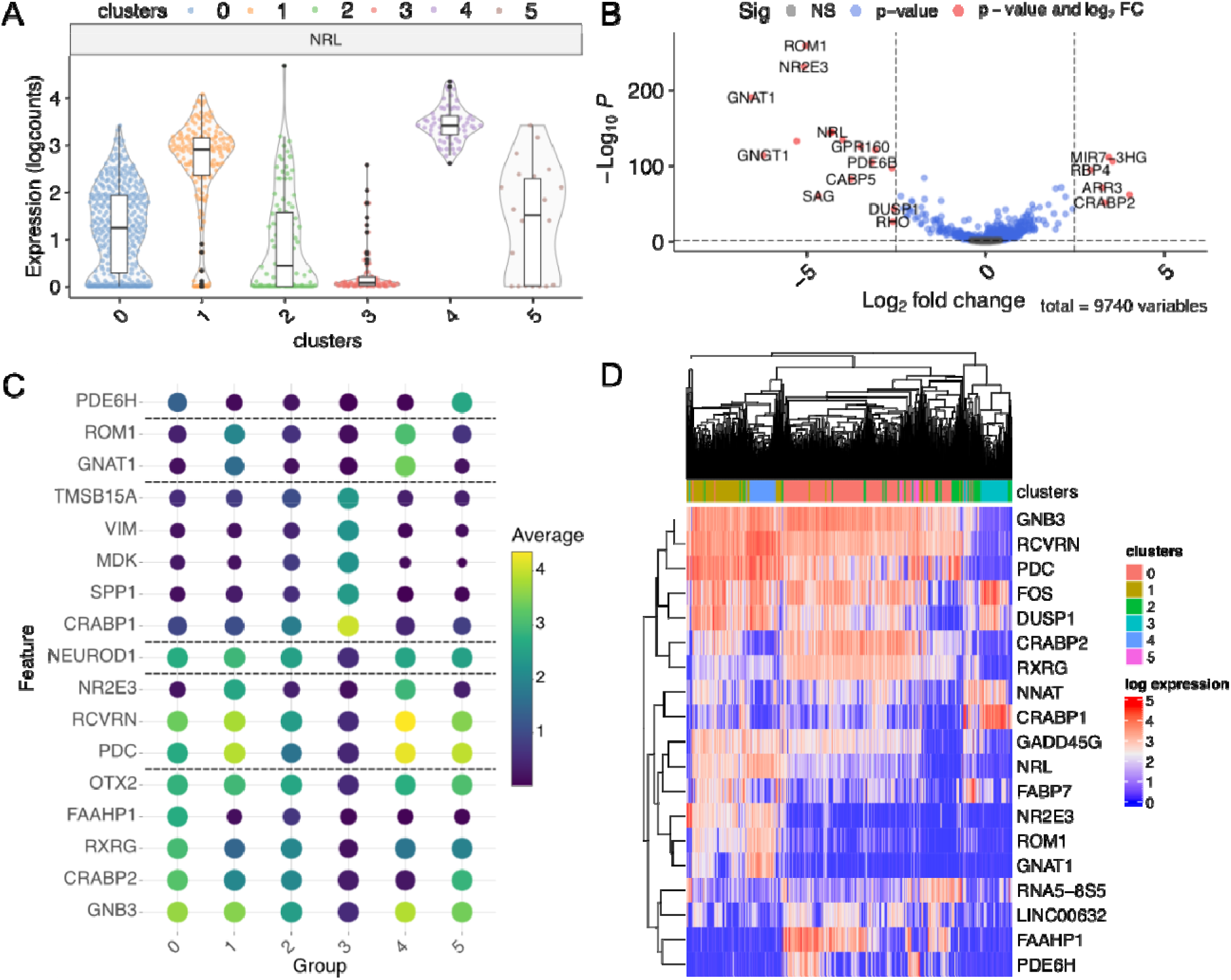
Gene expression plots. (A) Violin plot of expression colored by cell metadata. (B) Volcano plot after differential expression. (C) Cluster marker genes. (D) Heatmap of highly variable genes.

If the input SingleCellExperiment contains both gene and transcript counts, the Transcripts tab allows display of a gene’s constituent inferred transcripts as stacked bar plots for different groups (e.g., cluster or sample age) (Fig. 5A) or overlaid on PCA, tSNE, and UMAP embeddings (Fig. 5B, Supp. Fig. S1B). The Coverage tab allows plotting of exon read coverage with log or linear y-axis scales using the *wiggleplotr* R package [27] (Fig. 5C). All plots shown are interactive using plotly capabilities [28], which allows plots to be viewed at various zoom levels and allows cells to be identified by placing the cursor over dots (in embeddings) or over lines (in histograms). User- customized plots can be downloaded in raster (.png) or vector (.svg, .pdf) image formats for further analysis and presentation.

**Figure 5:**
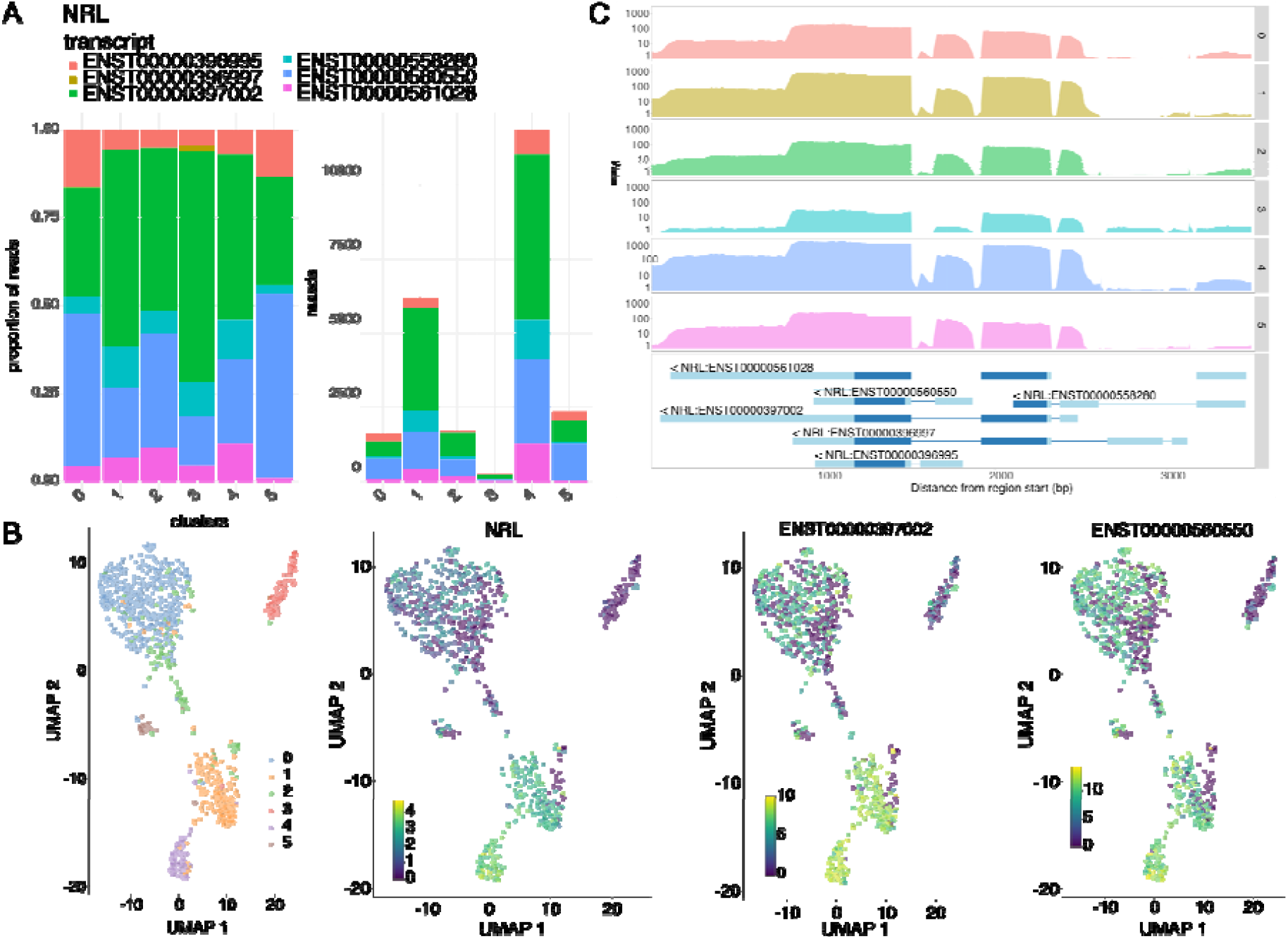
Transcript-specific plots. (A) Transcript composition by cell group as stacked bar plots. (B) *NRL* expression of constituent transcripts and aggregated gene expression. (C) Coverage of gene regions colored by cell metadata using wiggleplotr.

The Subset tab enables iterative analyses of data subsets defined by an uploaded delimited file or by using the cursor to select groups of cells from a dimensionally reduced plot. Subsetting of single or batch integrated data will trigger renewal of all relevant preprocessing steps including dimensional reduction, clustering, and marker genes, as well as integration based on a ‘batch’ variable. After subsetting, a reset option can be triggered within the subset tab to return data to an initial state. In the Regression tab, confounding cell-cycle effects can be regressed using cyclone [29]. The Formatting tab displays the original cell metadata and allows reformatting by in-app editing or by uploading a minimal file with cell identifiers and any new cell metadata to be added. Resulting reformatted tables can be exported as .csv files. All analysis steps including software versions are specified in a convenient Technical Information tab.

## Discussion

Chevreul is an open-source R package and R Shiny app that empowers researchers without prior programming experience to analyze scRNA-seq data. Chevreul allows processing, visualization, and interactive analysis in a standardized framework. The built-in Shiny app includes plotting functions specifically for full-length scRNA-seq data, including exon coverage and inferred isoform usage summarized according to the sample characteristics represented in the cell metadata. Inferring isoform usage, in turn, may reveal cell type-specific transcript isoform use and enable distinctions of closely related states not visible with gene expression alone [23]. The Chevreul application includes extensive guidance materials and allows users to visualize a wide range of parameters, enabling transparent and reproducible scRNA-seq analyses.

## Methods

Open-source software used in Chevreul package include:

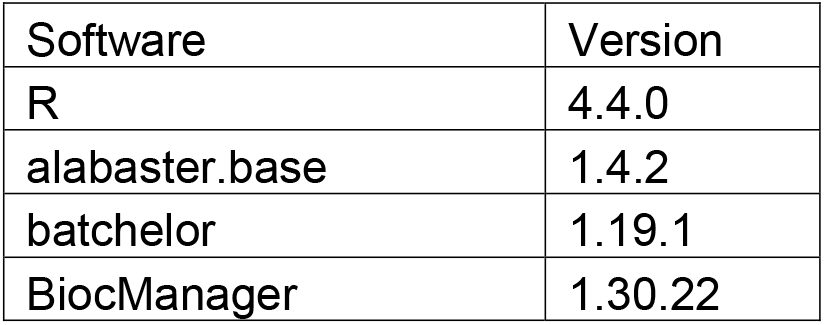
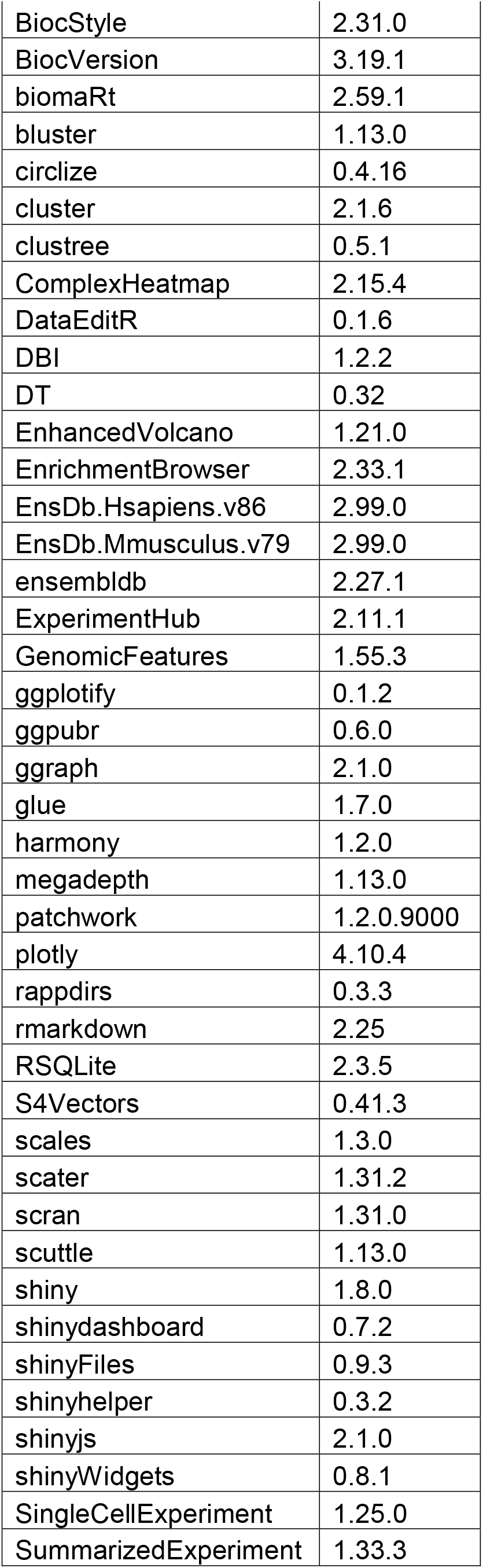
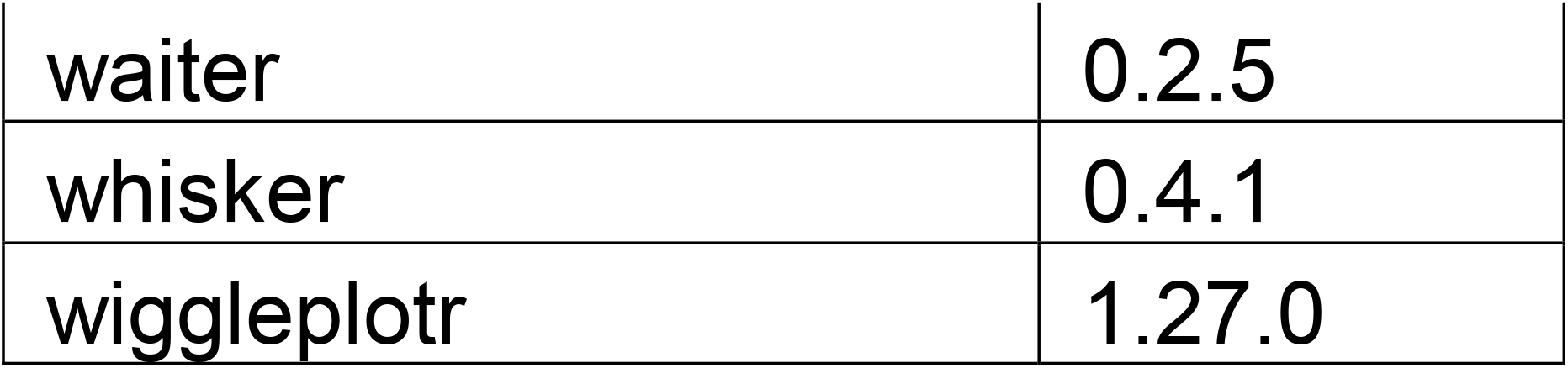

## Software availability

The Chevreul package is freely available at GitHub https://github.com/cobriniklab/chevreul. A companion package for processing and display of Seurat objects is available at https://github.com/cobriniklab/seuFLViz. Chevreul is registered in the bio.tools database at biotoolsID (biotools:chevreul) and the Scicrunch.org database at Research Resource Identification Initiative ID (RRID:SCR_026966).

### Summary information

Project name: Chevreul

Project home page: https://cobriniklab.github.io/chevreul/

Operating system(s): Platform independent

Programming language: R 4.4.0

License: MIT

### Availability of Test Data

A test datasetformatted as a SingleCellExperiment object can be found at https://github.com/cobriniklab/chevreuldata.

## Declarations

### List of abbreviations

PCA: principal component analysis
scRNA-seq: single cell
RNA: sequencing
tSNE: t-distributed stochastic neighbor embedding
UMI: unique molecular identifier
UMAP: Uniform Manifold approximation and projection

## Ethical Approval

Not applicable.

## Author’s Contributions

KS: Conceptualization, investigation, formal analysis, software, methodology, validation, data curation, writing – original draft preparation, writing – review and editing; visualization, project administration.

BB: Investigation, software, data curation, visualization.

DC: Supervision, project administration, writing - review & editing, funding acquisition.

---

**Supplementary Figure S1:**
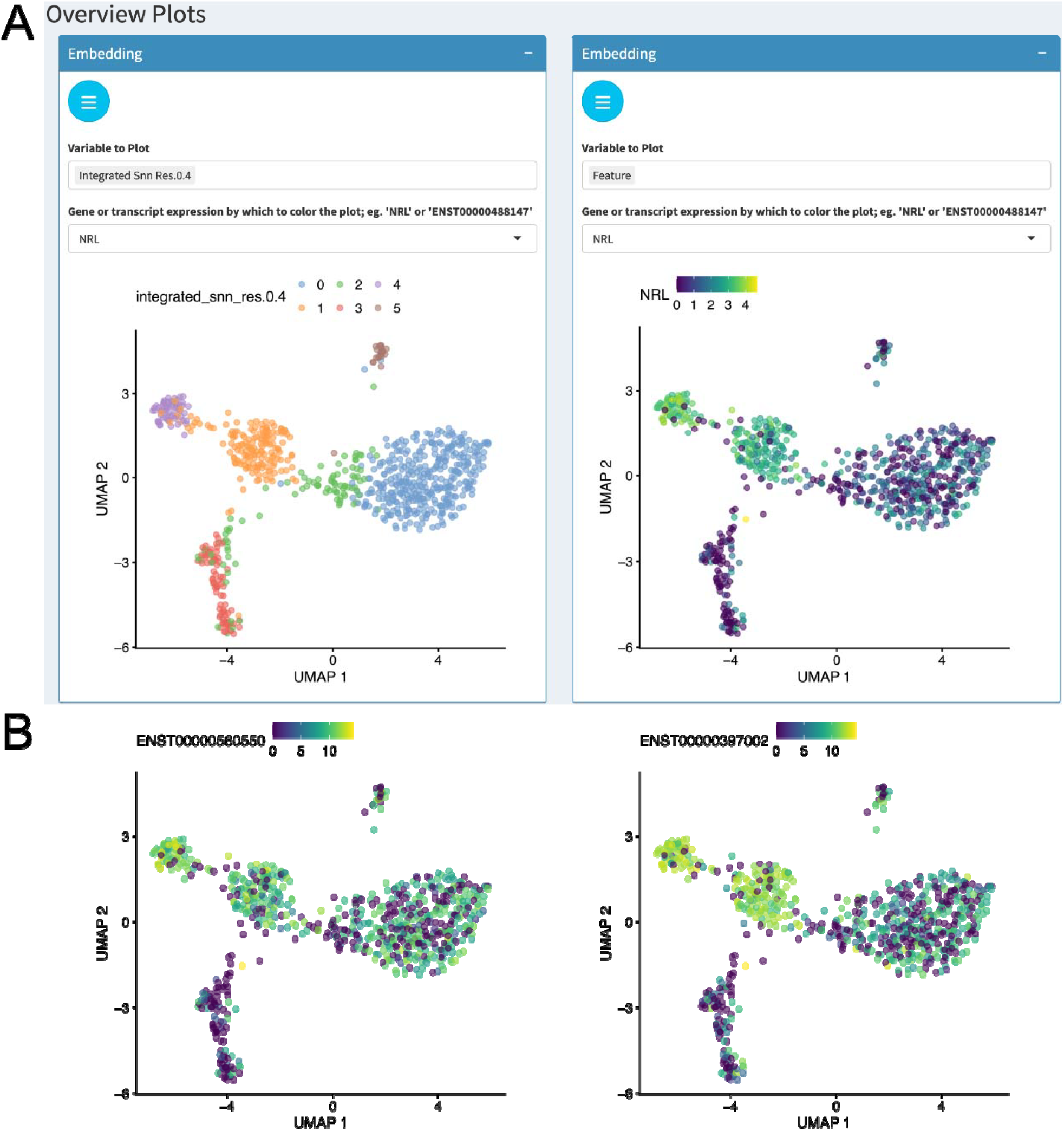
Analysis overview plots processed with Seurat. (A) UMAP embedding overlaid with metadata for cluster cell type (*left*) or expression of *NRL* (*right*). (B) Transcript-specific plots processed with Seurat. Expression of *NRL* transcripts ENST00000397002 and ENST00000560550 aggregated gene expression.

